# Alterations in Cell Motility, Proliferation, and Metabolism in Novel Models of Acquired Temozolomide Resistant Glioblastoma

**DOI:** 10.1101/232447

**Authors:** DM Tiek, JD Rone, GT Graham, EL Pannkuk, BR Haddad, RB Riggins

**Author notes:** Correspondence should be addressed to DM Tiek and RB Riggins. **Address**: Georgetown University, 3970 Reservoir Road NW, E412 Research Bldg., Washington, DC 20057.

## Abstract

Glioblastoma (GBM) is an aggressive and incurable tumor of the brain with limited treatment options. Current first-line standard of care is the DNA alkylating agent temozolomide (TMZ), but this treatment strategy adds only ~4 months to median survival due to the rapid development of resistance. While some mechanisms of TMZ resistance have been identified, they are not fully understood. There are few effective strategies to manage therapy resistant GBM, and we lack diverse preclinical models of acquired TMZ resistance in which to test therapeutic strategies on TMZ resistant GBM. In this study, we create and characterize two new GBM cell lines resistant to TMZ, based on the 8MGBA and 42MGBA cell lines. Analysis of the TMZ resistant (TMZres) variants in conjunction with their parental, sensitive cell lines shows that acquisition of TMZ resistance is accompanied by broad phenotypic changes, including increased proliferation, migration, chromosomal aberrations and secretion of cytosolic lipids. Importantly, each TMZ resistant model captures a different facet of the “go” (8MGBA-TMZres) or “grow” (42MGBA-TMZres) hypothesis of GBM behavior. These model systems will be important additions to the available tools for investigators seeking to define molecular mechanisms of acquired TMZ resistance.

## Introduction

Glioblastoma (GBM) is the most common brain cancer among adults and confers an abysmally low overall survival with only 5% of patients surviving at the 5-year mark^1^. Over the past 33 years – 1980-2013 – 570 clinical trials were conducted where almost 33,000 patients were treated with different novel therapeutics to better understand and treat GBM ^2^ From these extensive studies one chemotherapeutic agent – temozolomide (TMZ) – was found to moderately improve overall survival ^3^ In the last decade there has been little advancement in treatment, with the standard of care being radiotherapy and surgery, followed by TMZ^4^. However, resistance to TMZ is rapid, and a broadly effective second line of treatment has not yet been discovered^5^. For these reasons, we need better models to understand mechanisms of TMZ resistance and how to develop improved therapies for the future.

Cell line models have been invaluable in elucidating the molecular mechanisms behind the uncontrolled growth of cancer cells. As resistance to TMZ is rapid in clinical models, cell lines were used to better understand the mechanism behind the initial efficacy of TMZ sensitivity. TMZ is a prodrug that is preferentially activated in a more alkaline environment, which the brain provides, that spontaneously breaks down to highly reactive methyldiazonium cations. These byproducts preferentially methylate DNA bases at the *N7*-guanine, *N3*-adenine, and *O6*-guanine positions. The cytotoxic effect was shown to be mainly through the *O6*-guanine adduct, which can be reversed by the *O6*-methylguanine methyl transferase (MGMT). MGMT is a suicide repair protein which removes the *O6*-guanine adduct and allows for proper DNA repair^6^. Even with this information, targeting MGMT with both small molecule drugs and mimetics has been unsuccessful, and its expression does not always correlate with resistance to TMZ^7–11^. Therefore, new models of resistance need to be developed in order to better define molecular mechanisms of GBM resistance to TMZ.

In this study, we present two unique TMZ-resistant GBM cell lines derived from the 42MBGA and 8MBGA parental lines which were first characterized in 1997. The 42MBGA line was derived from a temporal lobe tumor resected from a 63-year old male. The 8MGBA line was derived from a frontal lobe tumor resected from a 54-year old female^12^. After continual exposure to TMZ in culture, both cell lines no longer undergo TMZ-mediated G2/M arrest. As the cell lines acquired resistance to TMZ both cell lines gained expression of MGMT, showed an increase in nuclear size and chromosome number, and an increase of early endosomes and extracellular lipids. However, as they differed in their growth and migratory capacity, these cell lines may give us insight into the “go or grow” model for GBM recurrence^13^. 42MGBA-TMZ resistant cell line had an increase in growth capturing the “go” of the “go or grow” model for resistance. By contrast, the 8MBGA-TMZ resistant cell line had an increase in migration encompassing the “grow” aspect of resistance. These models will be useful tools for the field to better define essential drug resistance mechanisms in GBM. These two TMZ-resistant (TMZres) models from both a male and female patient have evolved distinct resistant phenotypes and therefore serve as valuable resources to investigate the heterogeneity of TMZ-resistant GBM.

## Results

### Acquired temozolomide resistance decreases sensitivity to BCNU

To better recapitulate acute temozolomide (TMZ) resistance models, we created cell lines that were resistant to 200 μM TMZ, and were challenged in 100 μM TMZ for all experiments. The overall time to acquired resistance varied from approximately 2 months for the 8MBGA cell line to ~3 months for the 42MBGA cell line. Cells were defined as resistant when each no longer showed a G2/M arrest or apoptosis in response to TMZ treatment (Figure 1a-d). While both parental cell lines (WT) exhibited a significant increase in the SubG1, or apoptotic, fraction following TMZ treatment, this was lost in the TMZres variants (Figure 1e). However, the shape of the SubG1 curves differed between the two cell lines – 8MGBA-WT cells treated with TMZ showed a defined SubG1 peak typically associated with apoptosis, while 42MGBA-WT cells did not (Sup Figure 1a,b). This was corroborated by immunoblotting for cleaved poly (ADP-ribose) polymerase (PARP), which was observed in 8MGBA-WT but not 42MGBA-WT cells (Sup Figure 1c,d) Expression of the O6-methylguanine methyltransferase (MGMT) was detectable in both resistant cell lines, but absent in both parental lines (Figure 1f).

**Figure 1.**
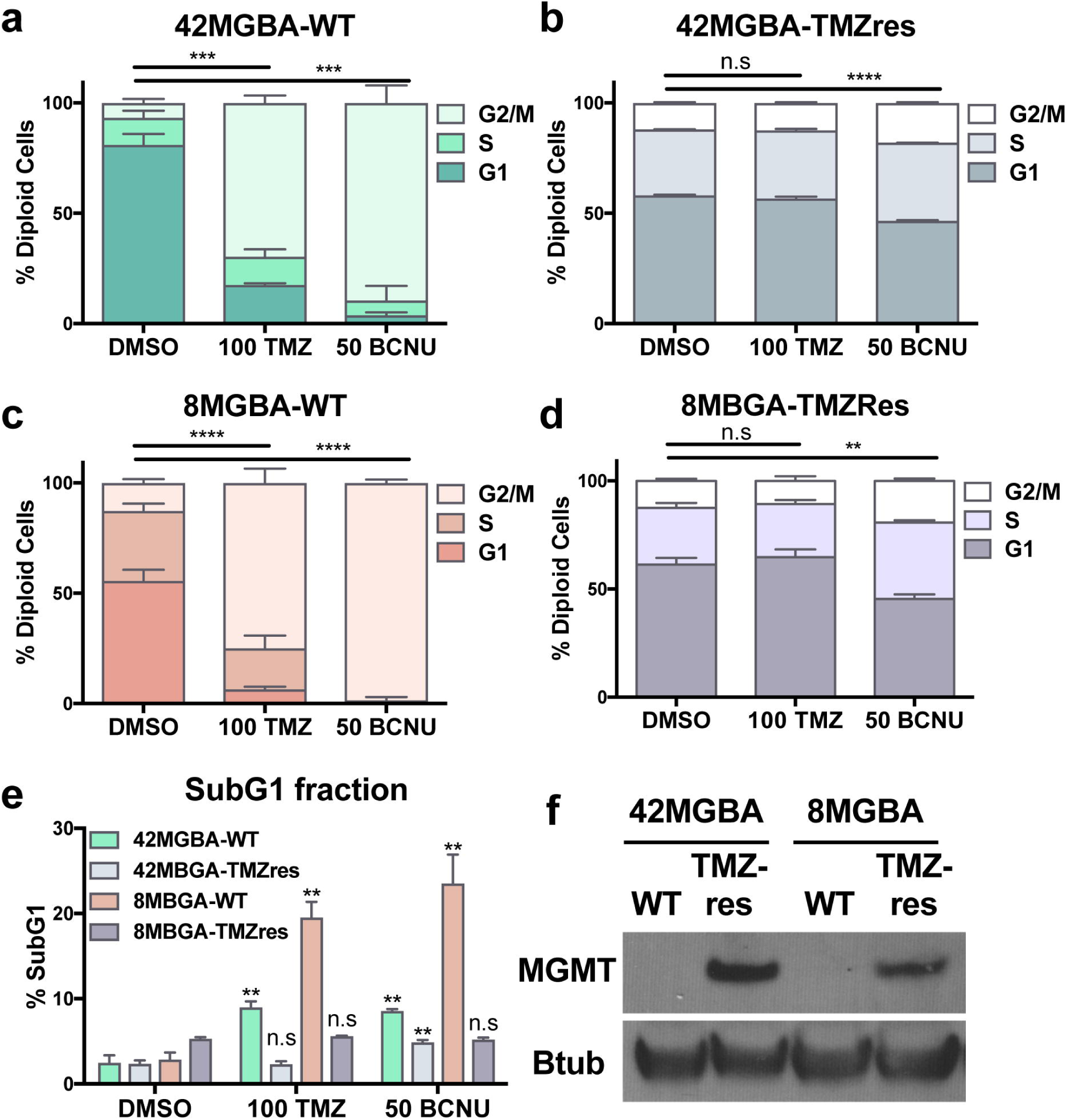
Acquired TMZ resistance. a,c) Cell cycle analysis of the parental 42MGBA and 8MGBA cell lines with 100 uM TMZ and 50 uM BCNU treatment for 72 hr. One-way ANOVA p = 0.0003; p = <0.0001 b,d) Cell cycle analysis of 42MGBA- and 8MBGA-TMZres cell lines with 100 uM TMZ and 50 uM BCNU treatment for 72 hours. One-way ANOVA p = <0.0001; p= 0.0010. e) Cell cycle analysis of SubG1 fraction for parental and resistant cell lines. t-test of treatment vs DMSO 42MGBA-WT p = 0.0046, 42MBGA-TMZres p = 0.0065, 8MBGA p = 0.0011. f. TMZres cells express MGMT.

As others have shown that acquired resistance to the previous standard of care for GBM, bis-chloroethylnitrosourea (BCNU), leads to an increased sensitivity to TMZ^14^, we tested whether our TMZ-res cells exhibited altered responsiveness to BCNU. Since the resistant cell lines are populations selected from the wild type cell lines they have a more reproducible phenotype as the variability or heterogeneity of the cell line has decreased. Therefore, the TMZ-res cells still showed a statistically significant G2/M arrest upon BCNU treatment. However the biological effect is dramatically reduced, with a 74% increase in G2/M vs 8% increase in G2/M for the 42MBGA-WT vs. 42MBGA- TMZres cells compared to controls, respectively (Figure 1a,b). This is similar to 8MGBA-WT and 8MGBA-TMZres cells, where the increase in G2/M fraction is 86% vs. 7% compared to controls, respectively (Figure 1c,d).

### Increase in nuclear size and chromosome number with TMZ resistance

The mechanism of action for TMZ is to induce DNA damage and give rise to cell cycle arrest in the G2/M phase^5^. As cells have already copied their DNA at this point in the cell cycle, if a cell is to survive TMZ-treatment they may also be retaining the extra copies of chromosomes that have already been duplicated. The 42MGBA-WT cell line has a hypertetraploid karyotype (88-95 chromosomes) with 8% polyploidy, while the 8MGBA-WT cell line has a hyperdiploid karyotype with 15% polyploidy and contains 47-52 chromosomes (DSMZ, https://www.dsmz.de/). 42MGBA-TMZres cells showed a significant increase in total nuclear area (Figure 2a), and their nuclear morphology became multi-lobed (Figure 2b). Total nuclear area was also significantly increased in 8MGBA-TMZres cells, though their nuclei retained an oblong or circular morphology (Figure 2a,b). Given the strikingly different nuclear morphology of the two resistant variants, we cultured both TMZres lines in the absence of TMZ for three weeks to test the stability of the resistant phenotype. After re-challenging with TMZ treatment for 72 hours, we observed no significant change in cell cycle profile or the SubG1 peak in the 42MBGA-TMZres cell line, while the 8MBGA-TMZres line underwent a slight G2/M arrest and showed a unique SubG1 peak in both untreated and treated conditions suggesting a less stable phenotype (Sup Figure 2a-d).

**Figure 2.**
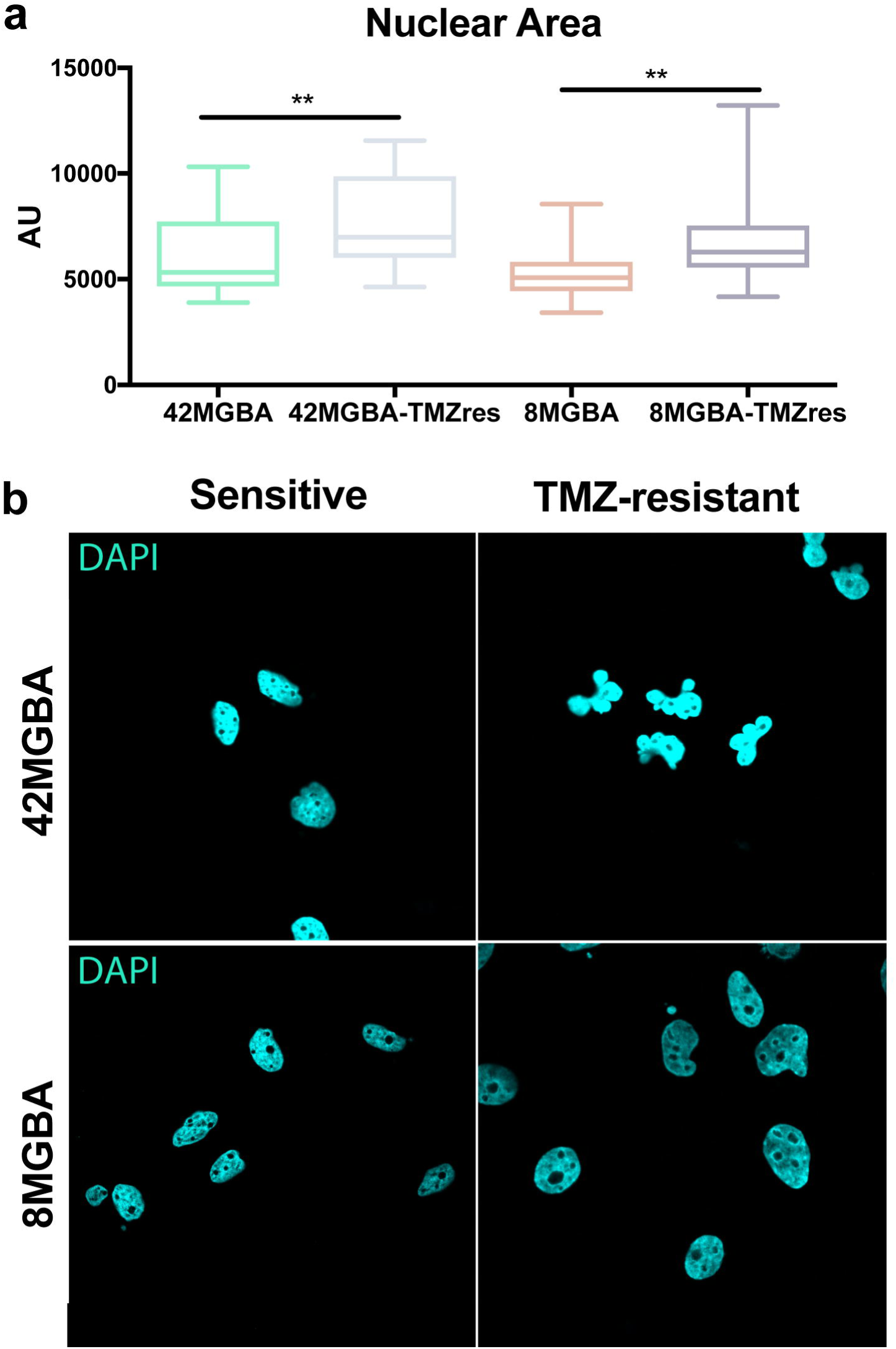
Changes in nuclear size and shape upon acquired TMZ resistance. a) Quantification of nuclear area by pixel size. AU = arbitrary units. Two-tailed t-test 42MBGA p = 0.0035; 8MBGA p = 0.0034. b) Representative image of nuclear size, quantified in a. Change in nuclear structure in 42MGBA-TMZres in comparison to 42MBGA-WT.

To determine whether increased nuclear size was associated with chromosomal gain, we assessed the overall chromosomal numbers in metaphase spreads from the TMZres cell lines compared to their respective parental lines and enumerated the copy number of chromosomes 17, in 8MGBA, and X, in 42MGBA, both in interphase and metaphase spreads from the parental cell lines and their resistant variants (Figure 3a). We saw an increase in overall chromosome number in the majority of the 42MBGA-TMZres and in a small subpopulation of 8MBGA-TMZres (Figure 3b). This again tracked with the stability of TMZ-resistance with the 42MBGA-TMZres showing a more stable phenotype. We then tested for TMZ-induced polyploidy in both resistant models by assessing the copy number of chromosome 17 in 8MBGA and its TMZres derivative, and of the X chromosome in 42MBGA and its TMZres derivative, by flueorescent in situ hybridization (FISH) in interphase nuclei. 42MBGA-WT showed two copies of the X chromosome in the majority of the nuclei (93%), and only 7% of the nuclei showed three or four copies. However, in the 42MBGA-TMZres, only 4% of the nuclei had 2 copies of the X chromosome and 96% had three to six copies. In contrast, while the majority of nuclei analyzed in the 8MBGA-WT cells showed two copies of chromosome 17 (94%), and 6% showed three or more copies, only 18% of the 8MGBA-TMZres cells had three or more copies of chromosome 17 (Figure 3c).

**Figure 3.**
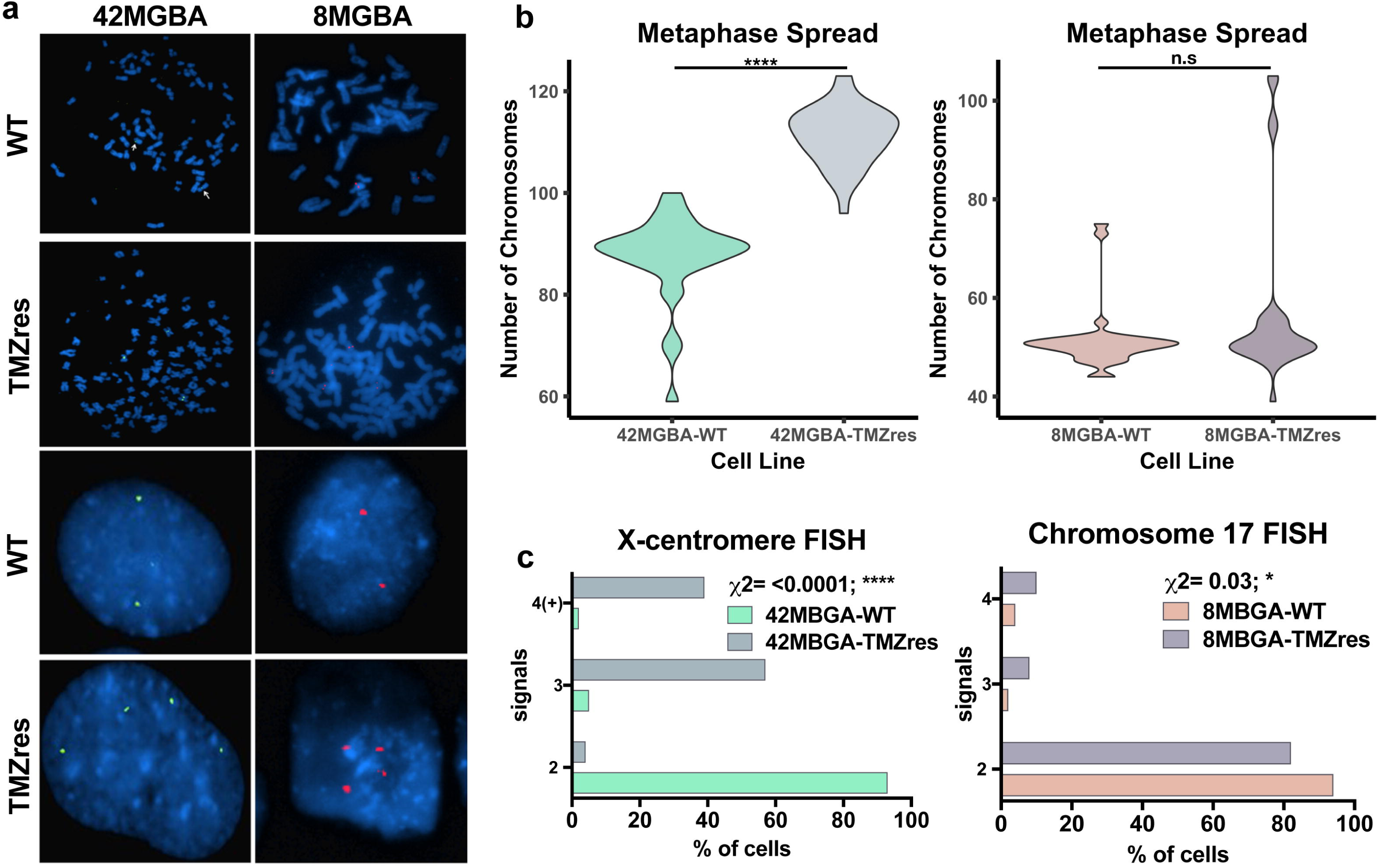
Acquired TMZ resistance increases chromosome number. a) Top 4 panels show metaphase spreads and chromosome number of sensitive and resistant 8MGBA and 42MGBA cell lines with probes targeting chromosomes 17 and X and the X, respectively. Bottom 4 panels show same probes as top 4 panels in interphase cells. b) Quantification of chromosomes from a, top 4 panels 42MGBA-WT vs –TMZres p = <0.0001. c) Quantification of probe signal from a, bottom 4 panels. Chi-squared test 8MBGA p = 0.03; 42MBGA p = <0.0001.

### Changes in proliferation, migration, and actin cytoskeleton

We then determined how TMZ-resistance affected cell size and proliferative vs. migratory phenotypes. 42MBGA-TMZres cell size was not changed vs 42MGBA-WT, though their basal growth rate was dramatically increased (Sup Figure 3a,b). They also showed a modest but nonsignificant reduction in cell migration (Figure 4a). In contrast, 8MBGA-TMZres cell size was significantly increased when compared to its parental cell line, while the basal growth rate was unchanged (Sup Figure 3a,b). 8MGBA-TMZres cells were significantly more migratory than 8MGBA-WT cells (Figure 4b). Enhanced cell migration correlated with increased F-actin stress fiber thickness in both TMZres models. There was no significant change in F-actin thickness in the 42MBGA-TMZres compared to 42MGBA-WT cells, while it was significantly increased in 8MGBA-TMZres when compared to 8MBGA-WT cells (Figure 4c,d).

**Figure 4.**
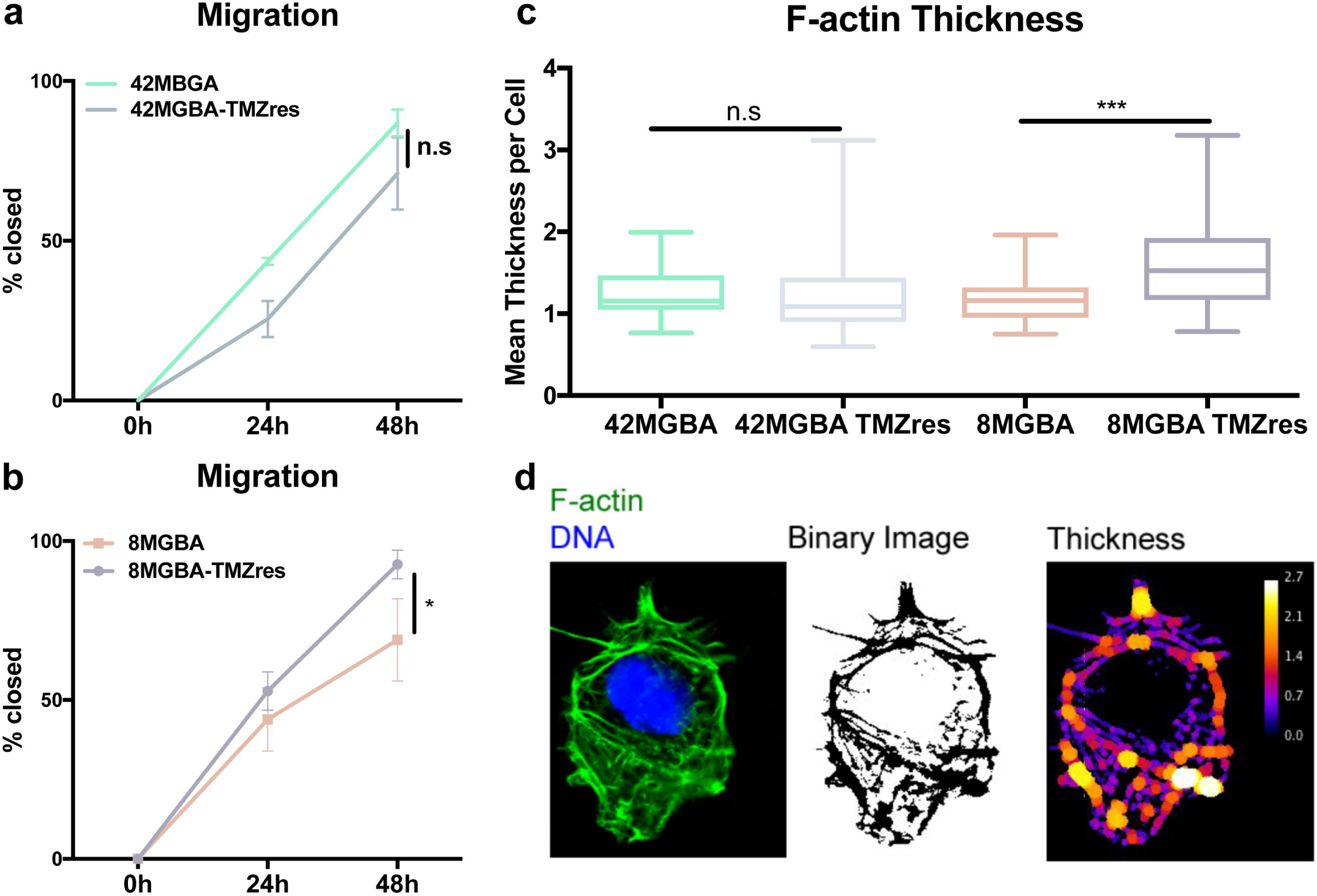
Changes in actin cytoskeleton and migration. a,b) Scratch-wound analysis for 2D migration over 48 hours. t-test at 48 hours 8MBGA p = 0.04 Mean thickness of F-actin filaments assessed by FIJI plug-ins as denoted in Methods section. Mann-Whitney U test 8MBGA p = 0.001. d) Representative image of quantification of c.

### TMZ resistance leads to an increase in intracellular vesicles and lipid metabolites

An advantage of selecting these resistant models was the ability to observe changes that occurred during and continued to persist after resistance was acquired. One observation was the number of intracellular puncta and extracellular vesicles that were expelled into the media as the cells became resistant to TMZ (Figure 5c). As there was no longer a dramatic increase in the SubG1 fraction (Figure 1), or cleaved PARP (Sup Figure 1c) upon resistance, we were able to rule out general apoptotic bodies. We therefore wanted to test if there was an increase in overall vesicle production in the TMZres vs WT cell lines. To accomplish this, we stained for the endosomal marker early endosome antigen 1 (EEA1). After analysis, we saw that there was a significant increase in the total number of endosomes in both TMZ-res cell lines compared to their parental lines (Figure 5a, b). Interestingly in two independent datasets – the Chinese Cancer Genomics Consortium (CCGC) and REpository for Molecular BRAin Neoplasia DaTa (REMBRANDT) – EEA1 expression increases from normal brain to GBM, and survival is significantly better for patients with tumors having lower expression of EEA1 (Sup Figure 4a-c). However, it was not only intracellular, or early endosome, vesicles that we observed to be different, but extracellular vesicles as well. As we were uncertain of the type of vesicles expelled into the media, we performed untargeted liquid chromatography and gas chromatography time-of-flight-based mass spectrometry (LC-MS, GC-ToF-MS) on cell culture media from the 8MGBA-WT and 8MBGA-TMZres cell line to identify differences in metabolites that may be associated with this phenotype. Putative results from the LC-MS showed an increase in the extracellular lipids LysoPC(16:0) and LysoPE(16:0) (Sup Figure 5). Interestingly, indoleacetaldehyde, a product of tryptophan metabolism, was also shown to be putatively increased in the 8MGBA-TMZres media. Tryptophan metabolism disruption has been shown previously as a feature of treatment adaptation in GBM cells^15^. Consistent with our observation of increased extracellular vesicles, there was an increase in palmitic acid and stearic acid lipids in the GC-TOF-MS metabolites from the 8MBGA-TMZres cell line conditioned media (Figure 6). The top hit from GC-TOF-MS was the non-aminogenic acid aminomalonate, an inhibitor of aminolevulinic acid synthase, the enzyme that catalyzes the rate limiting step of the protoporphyrin IX(PpI IX)/heme synthesis pathway^16^. This pathway is deregulated in many cancers, including GBM, and can be exploited to identify tumor area during surgery^17, 18^. The last significant metabolite was citric acid, which has been shown to decrease overall survival in GBM patients with higher citric acid in their cerebral spinal fluid^19^.

**Figure 5.**
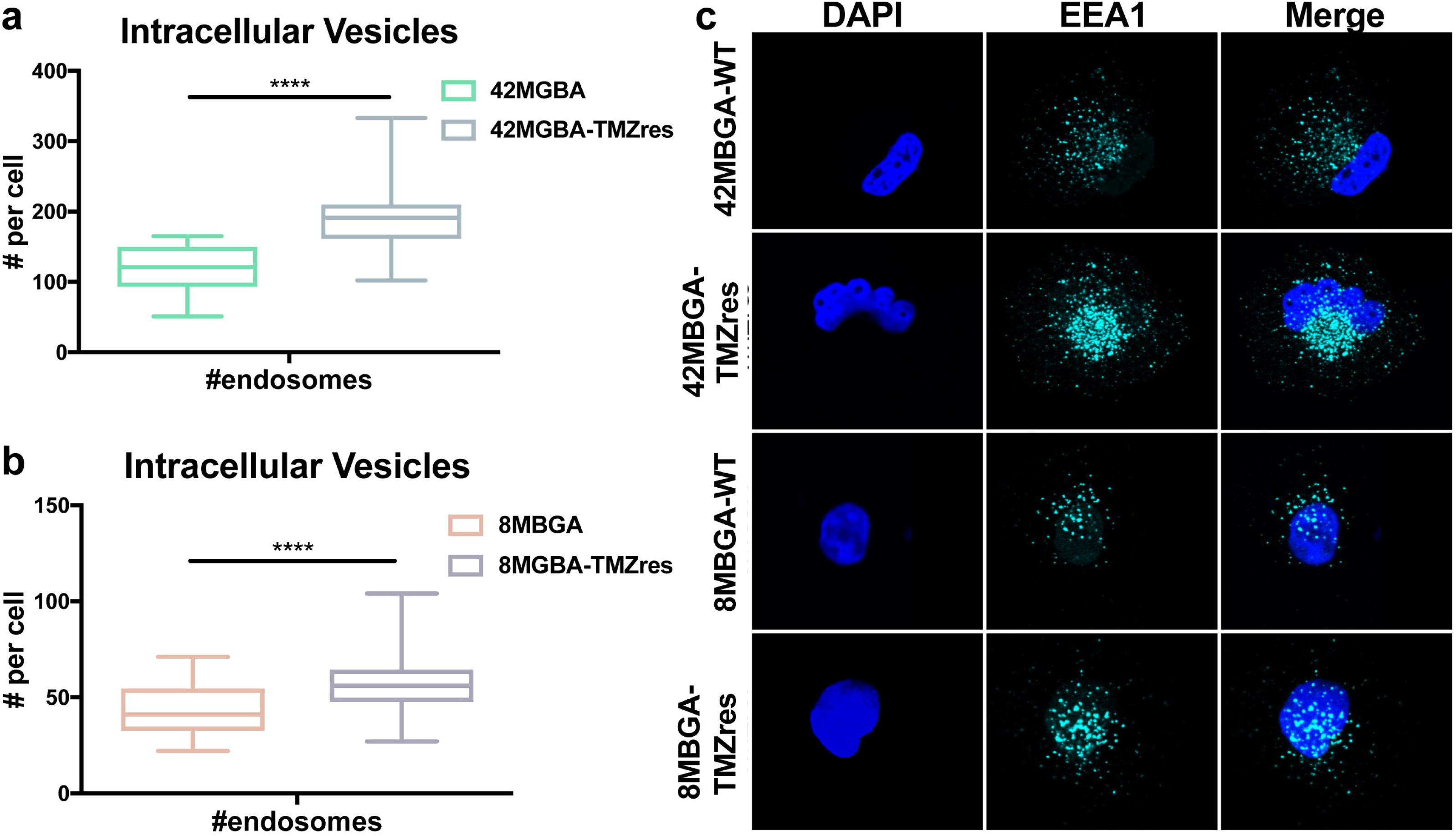
Increase of intracellular vesicles with TMZ resistance. a,b) Number of intracellular vesicles stained positive for the early endosomal marker EEA1 quantified between sensitive and resistant cell lines by immunofluorescence. t-test a) 42MGBA p = <0.0001; b) 8MBGA p = <0.0001. c) Representative image of EEA1 immunofluorescence.

**Figure 6.**
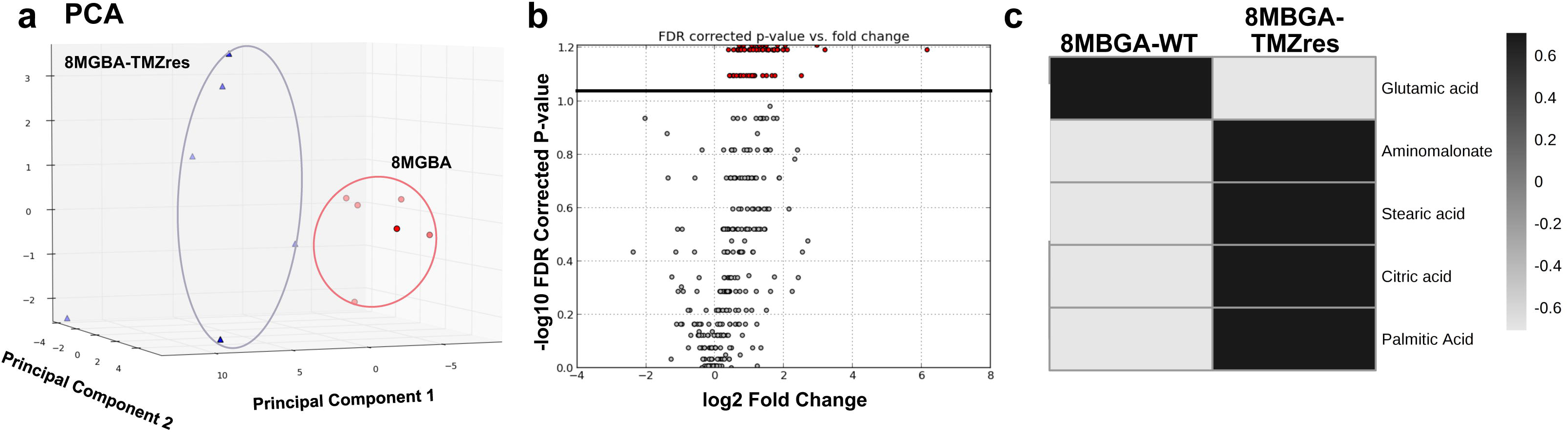
GC-MS analysis of 8MBGA-WT and –TMZres media. a) PCA scores plot of metabolites from 8MGBA and 8MGBA-TMZres cell culture media detected by Gc-MS and b) the corresponding volcano plot (significant ions above black line). Heatmap in c) shows significant metabolites between the groups (p = 0.005) and was validated through the retention index value and comparison of the electron impact spectra to the NIST 14 database. Log2 fold-change scale is shown.

## Discussion

In the present study, we created two new GBM cell lines that are resistant to the standard of care drug temozolomide (TMZ). True to the heterogeneous nature of clinical GBM, the phenotypes displayed by these resistant cells were similar, but not identical. TMZ treatment no longer induced G2/M arrest in either of these cell lines, and both were markedly less responsive to BCNU, the previous standard of care for GBM. Both resistant cell lines also gained expression of the MGMT protein, which is responsible for removing one of the DNA adducts caused by TMZ treatment^5^.

A key advantage provided by creating these resistant cell lines as opposed to isolation from a resistant tumor is the ability to directly compare them to their respective sensitive parental lines. There was an increase in nuclear size in both resistant cell lines when compared with their parental cell lines. In the 42MGBA-TMZres cell line there was also a change in nuclear morphology which has been described in other GBM cell lines that have acquired resistance to TMZ^20, 21^. This may be associated with stability of the resistant phenotype, as we observed that the 42MBGA-TMZres cell line resistant phenotype can be maintained without continuous TMZ treatment. However, the 8MGBA-TMZres cell line, which has not undergone nuclear structure reorganization, and showed two distinct populations with respect to metaphase spreads, regained partial sensitivity when cultured in the absence of TMZ.

Increases in nuclear size can be driven by increased chromosome numbers or impaired transport of mRNA or proteins out of the nucleus^22^. In these novel TMZ resistant models, we hypothesized that the increase in nuclear size was a function of retained chromosomes. This is consistent with the mechanism of action of TMZ, which induces a G2/M arrest in cells. Cells which escape TMZ-mediated cell cycle arrest and death have already replicated their DNA, and may therefore retain sister chromatids that were not able to separate during anaphase. Because both parental GBM cell lines were initially polyploid with distinct karyotypes, we assessed two different chromosomes – 17 for the 8MBGA-WT and –TMZres cell lines, and X for the 42MBGA-WT and –TMZres cell lines – as an indicator for chromosomal copy number alterations. Both TMZ resistant variants showed significant copy number gains. Of note, our chromosome 17 locus specific probe targets the STaT5 gene, which has already been implicated as a pro-tumorigenic factor in GBM^23^. The increased copy number of the *STAT5* locus in the 8MBGA-TMZres cell line is consistent with previous work and suggests that the observed increase in nuclear size is at least partially attributable to an increase in chromosome number. Because chromosome 17 existed in multiple copies in the 42MBGA-WT cell line, we chose an X-centromeric probe to test chromosomal copy number changes in the 42MBGA-TMZres line. Again, we saw an increase in X-chromosome copy number in the 42MBGA-TMZres line, and an overall increase in the total number of chromosomes. While it can be energetically disadvantageous to increase ploidy, we hypothesize that, as for STAT5, there may be X chromosome genes that when duplicated give a proliferative advantage to these cells.

Two major challenges in the clinical management of GBM, particularly after TMZ resistance has developed, are rapid tumor cell proliferation and widespread local migration or invasion^24^. However, these phenotypes are rarely displayed by the same cells, giving rise to the "go or grow” hypothesis to explain aggressive behavior and drug resistance^28^. Our novel models of TMZ-resistant GBM provide examples of both phenotypes. The 42MBGA-TMZres cell line did not exhibit any significant changes in migration, cytoskeletal structure, or cell size when compared to its parental cell line, but it did proliferate significantly faster. In contrast, the 8MBGA-TMZres cell line had a significantly increased migratory capacity, F-actin thickness, and overall cell size relative to 8MGBA-WT cells, but no accompanying increase in proliferation. These divergent alterations in basal growth rate and migratory capacity provide two unique models – one adapting the “go” (8MGBA-TMZres) and the other the “grow” (42MBGA-TMZres) strategy of TMZ resistance^13^. These four cell lines are therefore valuable tools that can be used to provide important insight into the differential mechanisms of TMZ resistance.

Acquisition of TMZ resistance was associated with a marked increase in the number of intracellular and extracellular vesicles in our models. The increase in early endosome marker EEA1 expression is interesting, as others have shown EEA1 to interact directly with RRAD, STAT3, and EGFR to regulate EGFR’s subcellular localization and resistance to TMZ ^25^ Clinically, higher EEA1 mRNA expression is associated with poor survival. EGFR and its structural variants have been targeted extensively in GBM, but with limited success. We suggest further study of the mechanisms by which EEA1 contributes to the TMZ resistant phenotype could provide a new way to target EGFR by modifying its subcellular localization rather than activation^26^. In addition to an accumulation of early endosomes, we observed vesicles expelled into the media of the resistant cell lines. Others have shown that TMZ-resistant glioma stem cells release more exosomes than their sensitive counterparts, which was consistent with our created models^27^. This was reflected in the putative increase of lipids from the LC-MS analysis. Another hit, indoleacetaldehyde, is a product of tryptophan metabolism, hyperactivation of which occurs in GBM and is thought to suppress anti-tumor immunity^15^. Extracellular lipids, stearic and palmitic acid, were also found and validated in the GC-TOF-MS analysis along with citric acid. Citric acid is a key regulator of metabolism, and when increased in the cerebral spinal fluid of GBM patients confers a lower overall survival^19^. The top hit from GC-TOF-MS was the non-aminogenic amino acid aminomalonate. Aminomalonate inhibits aminolevulinic acid synthase, the rate-limiting enzyme of the protophorphyrin IX (PpI IX)/heme synthesis pathway^16^. PpI IX pathway activity is exploited for intra-operative imaging in GBM, and has more recently been used for potential therapeutic gain^28, 29^. Defining the drivers of aminomalonate secretion in our TMZ-resistant models may be an important future strategy to identify when PpI IX pathway-based imaging and therapeutic approaches may be warranted.

In conclusion, we present two novel TMZ-resistant GBM cell lines from both a male and female patient that have distinct proliferative and migratory capacities while sharing certain GBM resistance characteristics such as increased chromosome number, nuclear size, and vesicle formation (Figure 7). These models will be useful tools for the field to better define essential drug resistance mechanisms and therapies in TMZ-resistant GBM.

**Figure 7.**
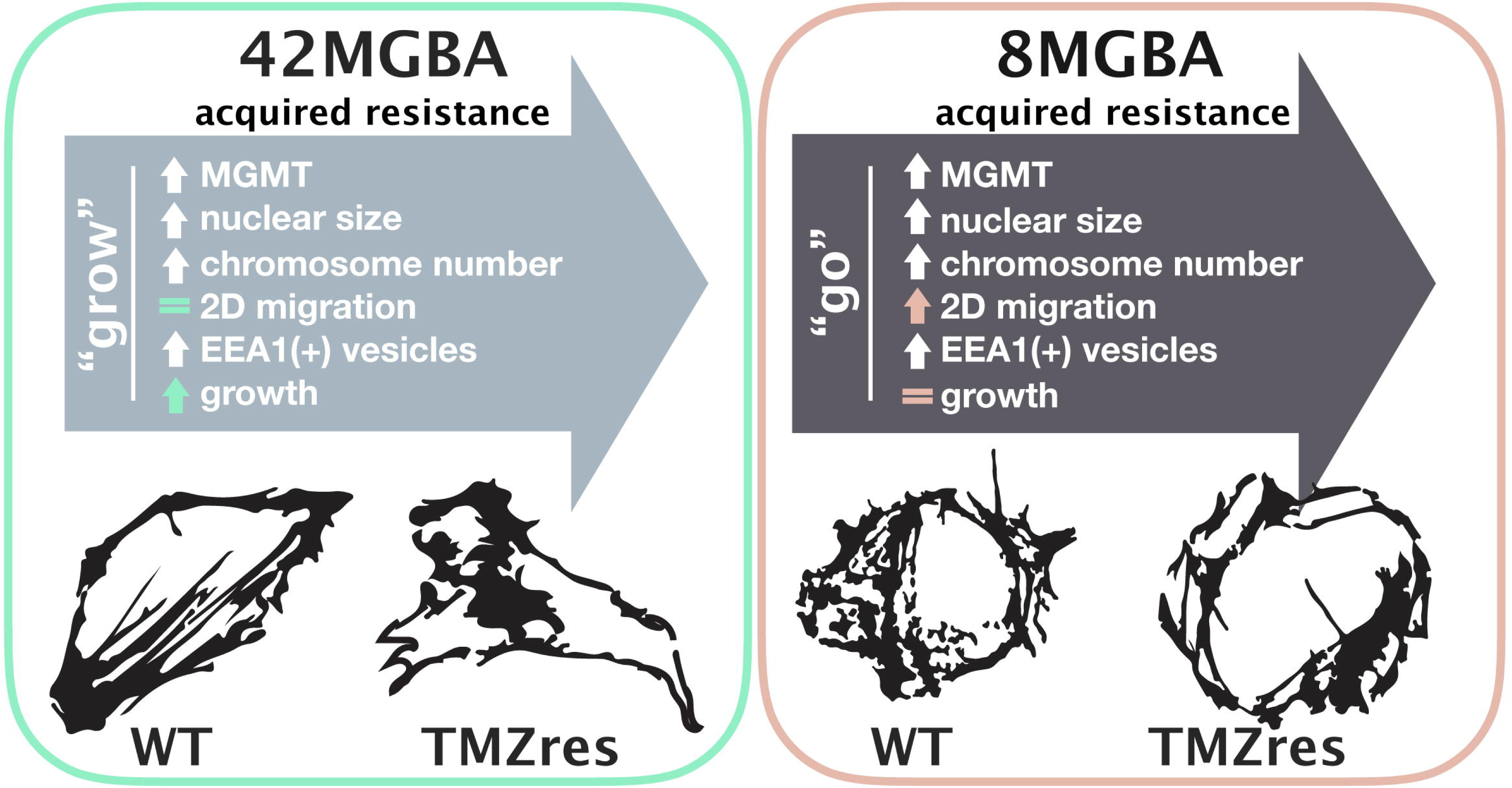
Summary figure of changes between wild-type (WT) cell lines upon acquired TMZ resistance (TMZres).

## Methods

### Cell Lines and Culturing Conditions

42MBGA and 8MGBA cell lines were provided by Dr. Jeffrey Toretsky (Lombardi Comprehensive Cancer Center (LCCC), Georgetown University, Washington DC). 42MBGA-TMZres and 8MGBA-TMZres were developed by our lab with constant exposure of the parental cell lines to increasing concentrations of TMZ, from 12.5, 25, 50, 100, 200 μM, over the course of 2-3 months. All cells tested negative for *Mycoplasma* contamination, and were maintained in a humidified incubator with 95% air: 5% carbon dioxide. All cell lines were fingerprinted by the LCCC Tissue Culture Shared Resource to verify their authenticity using the standard 9 STR loci and Y-specific amelogenin. Both the 42MBGA-TMZres and 8MGBA-TMZres fingerprinted the same as their parental cell line. 42MGBA and 8MBGA cells were grown in DMEM with 10% FBS. 42MBGA-TMZres and 8MBGA-TMZres cells were grown in DMEM with 10% FBS and 100 μM TMZ. TMZ (Selleckchem, Catalog No. S1237) was dissolved in DMSO to 130 mM and used at concentrations indicated. Bis-chloroethylnitrosourea (BCNU) was a kind gift from Dr. Esther Chang, and was dissolved in ethanol to 100mM before use at concentrations indicated.

### Cell Cycle Analysis

On day 0, cells were seeded at 100,000 – 150,000 cells per well in 6-well plastic tissue culture dishes one day prior to treatment with the indicated concentrations of drug. For experiments with TMZ or BCNU cells were treated for 72 hours. After 72 hour treatments, cells were collected, ethanol-fixed, stained with propidium iodide, and analyzed for cell subG1 (fragmented) DNA content and cell cycle profile by fluorescence activated cell sorting.

### Western Blotting

Cells were lysed in RIPA buffer for protein extractions and separated by a 4-12% gradient gel (Novex by Life Tech, NP0321BOX). They were then transferred onto Nitrocellulose membranes (Invitrogen, IB23001) with the iBlot 2 and probed with antibodies against MGMT (Cell Signaling, 2739S) and Beta-Tubulin (Sigma Aldrich, T7816). Proteins were detected with horseradish peroxidase-conjugated secondary antibodies (1:5000, GE Healthcare Life Sciences) and enhanced chemiluminescent detection HyGLO Quick Spray Chemiluminescent (Denville) for dark room development.

### Evaluation of chromosomal copy number changes induced by TMZ treatment

In order to assess the chromosomal copy number changes in the TMZ resistant cell lines compared to their respective parental lines, we evaluated the overall chromosomal copy number in metaphase spreads from each line and we also assessed copy number changes of a representative chromosome (chromosome 17 for 8MGBA and X for 42MGBA) in interphase nuclei, using fluorescence *in situ* hybridization (FISH) with chromosome specific probes. The choice of the two representative chromosomes (17 and X) was made based on reported karyotype analysis of the 2 parental cell lines showing a mostly diploid count for chromosome 17 in 8MGBA line and X in the 42MGBA (DSMZ, https://www.dsmz.de/). Chromosomes and interphase slide preparation was done using a standard protocol^30^. For metaphase chromosomal copy number determination, chromosomes were stained with 4’,6-diamidino-2-phenylindole (DAPI). For chromosome 17 enumeration in interphase nuclei, a 17q21.1-21.2 locus-specific probe (Stat5A/B locus) was used as described earlier^31^. For the X chromosome enumeration, we used an X chromosome centromeric probe obtained from Empire Genomics (Buffalo, NY). FISH analysis was performed using a standard protocol^31–33^.

### Scratch Wound Assays

Cells were plated at 150,000-200,000 cells/well, and allowed 48 hours to create a monolayer. After monolayer formation, a P200 tip was used to make the scratch. Images were taken at 0 hr, 24 hr, and 48 hr time points. Analysis was done in ImageJ^34^ determine percent closed with 0% being at 0hr.

### F-actin measurements

FIJI (ImageJ, ^34^) was used to measure the thickness of F-actin fibers. Grayscale images of phalloidin-stained cells were converted to binary using the Threshold tool (black/white). The BoneJ plugin ^35^ was then used to measure fiber thickness in pixels. The mean thickness per cell is reported for 28-29 individual cells per cell line.

### Immunofluorescent Staining

8MGBA-WT and 8MGBA-TMZres cells were seeded at a density of 50,000 cells onto 18mm coverslips in 12-well dishes. On the following day, the media was removed and cells were fixed and permeabilized in 3.2% paraformaldehyde (PFA) with 0.2% Triton X-100 in PBS for 5 minutes at room temperature. Three washes were performed with PBS in the 12-well plate, then coverslips were inverted onto 100 μl of primary antibody in the antibody block (0.1% gelatin with 10% normal donkey serum in PBS) on strips of parafilm and incubated for one hour. Coverslips were first stained with EEA1 (Cell Signaling, 3288), an early endosomal marker (1:100 dilution, 1 hr). Phalloidin (polymerized F-actin, Invitrogen, A12379, 1:200 dilution), and DAPI (DNA, 1:500 dilution) stains were added to the secondary antibody mixtures. Each coverslip was washed three times with PBS after primary incubation and then were inverted onto 100 μL of the appropriate secondary antibody, DAPI dihydrochloride, and (where appropriate) ActiStain-488-phalloidin (Cytoskeleton, Denver, CO) in antibody block in the dark for 20 minutes. Coverslips were again washed 3x with PBS, then gently dipped three times into molecular biology-grade water before inversion onto one drop of FLUOROGEL (Electron Microscopy Sciences, Hatfield, PA) then allowed to air-dry in the dark for at least 10 minutes. Slides were stored at 4°C until image collection on the LCCC Microscopy & Imaging Shared Resource’s Leica SP8 microscope with the 63X oil objective.

### Metabolomics

#### Ultra-performance liquid chromatography (UPLC) quadrupole time-of-flight (QTOF) mass spectrometry (MS)

Cell culture media (500 μL) was concentrated to 25 μL under N_2_. The 25 μL sample was deproteinized using cold acetonitrile (40%): methanol (25%): water (35%) (175 μL) containing internal standards (2 μM debrisoquine sulfate, 30 μM 4-nitrobenzoic acid). All solvents used were LC-MS grade (Fisher Scientific, Hanover Park, IL). Samples were vortexed and incubated on ice for 10 min. Incubated samples were centrifuged at 13000 rpm for 20 min at 4 °C. The supernatant was evaporated in a speed-vac with no heat. Dried samples were kept at −80 °C until analysis. For analysis, the dried samples were reconstituted in acetonitrile (1%): methanol (5%): water (94%) (200 μL). Samples were centrifuged for 20 min at 4 °C and the supernatant transferred to a vial for LC-MS analysis. Briefly, samples were injected (2 μL) into a Waters Acquity UPLC system coupled to a Xevo^®^ G2 QTOF-MS using Leucine enkephalin as Lockspray^®^ for accurate mass calibration. Data was processed, aligned, and analyzed using MassLynx^TM^ (Waters, Milford, MA), XCMS online^36^, and R^37^.

#### Gas chromatography time-of-flight mass spectrometry (GC-TOF-MS)

Internal standards (1 μM 4-nitrobenzoic acid) were added to culture media (500 μL) that was deproteinized as above and then evaporated in a speed-vac and transferred to a GC vial with a silanized insert. Derivatization and injection was performed with a Gerstel (Linthicum, MD) autosampler with automated liner exchange and cooled injection system. The dry residue was derivatized with 20 μL O-Methoxyamine-Hydrochloride (MOX; Thermo Scientific prod # TS-45950) at 40 °C with constant agitation (30 min), and then with 80 μL of N-Methyl-N-(trimethylsilyl) trifluoroacetamide (MSTFA) / with 1% trimethychlorosilane (TMCS) (Thermo Scientific prod # TS48915) at 40 °C with constant agitation (30 min). Samples were incubated at 20 °C for 4 hrs before injection (1.5 μL) for GC-MS analysis.

Samples were injected in split mode (1:5) into an Agilent 7890B GC system (Santa Clara, CA, USA) that was coupled to a Pegasus HT TOF-MS (LeCO Corporation, St. Joseph, MI, USA). Separation was achieved on a Rtx-5 w/Integra-Guard capillary column (5% diphenyl / 95% dimethyl polysiloxane; 30 m × 0.25 mm ID, 0.25 μm film thickness; Restek Corporation, Bellefonte, PA, USA), with helium as the carrier gas at a constant flow rate of 1.0 mL/min. The temperature of the inlet, transfer line, and ion source was set to 150, 270, and 230 °C, respectively. The GC temperature programming was set to 0.2 min of isothermal heating at 70 °C, followed by 10 °C/min ramp to 270 °C, a 4.0 min isothermal heating of 270 °C, 20 °C/min ramp to 320 °C, and a 2.0 min isothermal heating of 320 °C as previously described^38^. Electron impact ionization (70 eV) at full scan mode (*m/z* 40–600) was used, with an acquisition rate of 30 spectra/sec. Fatty acid methyl esters (C_4_-C_24_ FAMEs; Sigma-Aldrich), alkanes (alkane standard mix [C_10_-C_40_]; Sigma-Aldrich), and quality control standards (oxalic acid, malonic acid, malic acid, citric acid, methionine, and 2-butenedioic acid) were run to ensure reproducibility of retention indices and derivatization. Data was processed, aligned, and analyzed using ChromaTOF v. 4.51.6.0 with the statistical compare function (Leco, St. Joseph, MI) and in house software MetaboLyzer^39^ as previously described^38^.

### Crystal violet assay

Cells were seeded at a density of 1,000 cells per well in 3, 96-well plastic tissue culture dishes per cell line on day 0. On day 1, one plate was stained with crystal violet (Sigma,C0775). For staining, plates were rinsed 1 time with 1X PBS to remove excess cellular debris. After, 100 μL of 3.2% PFA was added to each well and incubated at room temperature for 5 minutes followed by three washes in 1X PBS. Then, 200 μL of 0.5% crystal violet in 25% methanol was added to each well and incubated at 4C for 10 min. The stain was then removed and the plate was rinsed 4–6X with diH2O to remove excess stain. The plates were left to air-dry overnight. On day 8, all plates were rehydrated with 100% methanol and read at an absorbance of 550 nm.

### Cell size quantification

Cell size was determined by the Countess 2 (Invitrogen) using trypan blue dye exclusion of dead cells.

### Statistical Analysis

Results are represented as mean values ± standard error (SD) and considered statistically significant where p-value <0.05. Three independent biological replicates were done for each experiment, with the exception of metabolomics where six independent replicates were performed. Statistical significance has been calculated using GraphPad Prism version 7.00 for Mac, GraphPad Software, La Jolla California USA (www.graphpad.com) for the Student’s t-test, Mann Whitney U test, one-way ANOVA with Dunnett’s correction, or χ2 analysis of 2×3 contingency table. The RStudio packages survival, GGally, ggplot2, readr, and magrittr were used for violin plots and big data sets (CCGA, REMBRANDT)^37^.

## Acknowledgements

We wish to thank Drs. Maria Laura Avantaggiati, Esther Chang, Amrita Cheema, Kirandeep Gil, Steven Peyton, Karen Creswell, Dan Xun, Jeffrey Toretsky, and Todd Waldman for sharing reagents, scientific insights, technical assistance, and/or editorial comments on the manuscript.

## Author Contribution Statement

DMT contributed to study design, performed experiments, analyzed data, and wrote the paper. JDR performed experiments. GTG analyzed data. eLp analyzed data. BRH contributed to study design, and analyzed data. RBR contributed to study design, analyzed data, and wrote the paper. All authors reviewed, edited, and approved the manuscript.

## References

1. Yang L-J, Zhou C-F, Lin Z-X. Temozolomide and Radiotherapy for Newly Diagnosed Glioblastoma Multiforme: A Systematic Review. Cancer Investigation. 2013;32(2):31–36. doi: 10.3109/07357907.2013.861474.

2. Stuart A Grossman SGE. Published glioblastoma clinical trials from 1980 to 2013: Lessons from the past and for the future. doi: 10.1200/JC0.2016.34.15_suppl.e13522#.

3. R S, WP M, van den Bent MJ, et al. Radiotherapy plus Concomitant and Adjuvant Temozolomide for Glioblastoma. New England Journal of Medicine. February 2005:987–996.

4. Mirimanoff R-O, Gorlia T, Mason W, et al. Radiotherapy and Temozolomide for Newly Diagnosed Glioblastoma: Recursive Partitioning Analysis of the EORTC 26981/22981-NCIC CE3 Phase III Randomized Trial. Journal of Clinical Oncology. 2016;24(16):2563–2569. doi:10.1200/JC0.2005.04.5963.

5. Lee SY. Temozolomide resistance in glioblastoma multiforme. Genes & Diseases. 2016;3(3): 198–210. doi: 10.1016/j.gendis.2016.04.007.

6. wasim. Temozolomide: Mechanisms of Action, Repair and Resistance. December 2011:1–13.

7. Zhang K, Wang X-Q, Zhou B, Zhang L. The prognostic value of MGMT promoter methylation in Glioblastoma multiforme: a meta-analysis. Familial Cancer. 2013;12(3):449–458. doi: 10.1007/s10689-013-9607-1.

8. Binabaj MM, Bahrami A, ShahidSales S, et al. The prognostic value of MGMT promoter methylation in glioblastoma: A meta-analysis of clinical trials. Journal of Cellular Physiology. 2017;233(1):378–386. doi: 10.1002/jcp.25896.

9. Yin A-A, Zhang L-H, Cheng J-X, et al. The Predictive but Not Prognostic Value of MGMT Promoter Methylation Status in Elderly Glioblastoma Patients: A MetaAnalysis. Lim M, ed. PLoS ONE. 2014;9(1):e85102. doi: 10.1371/journal.pone.0085102.

10. Czeremszyńska B, Bujko M, Ibron G, Onap-Karnak A, Nawrocki S. Treatment results, including clinical prognostic factors and MGMT gene promoter methylation, in patients with glioblastoma multiforme in Warmia and Mazury Oncology Centre. Wspófczesna Onkologia. 2011;4:198–202. doi:10.5114/wo.2011.24313.

11. D’Alessandris QG. Prognostic Impact of MGMT Promoter Methylation in Glioblastoma - A Systematic Review. Journal of Cancer Science & Therapy. 2014;06(04). doi: 10.4172/1948-5956.1000261.

12. A PERZEDDVA IMPMIBJS. Characterization of two new permanent glioma cell lines 8-MG-BA and 42-MG-BA*. 1998;45(1):25–29.

13. Hatzikirou H, Basanta D, Simon M, Schaller K, Deutsch A. “Go or grow”: the key to the emergence of invasion in tumour progression? Math Med Biol. 2012;29(1):49–65. doi: 10.1093/imammb/dqq011.

14. Yamauchi T, Ogawa M, Ueda T. Carmustine-resistant cancer cells are sensitized to temozolomide as a result of enhanced mismatch repair during the development of carmustine resistance. Mol Pharmacol. 2008;74(1):82–91. doi: 10.1124/mol.107.041988.

15. Sordillo PP, Sordillo LA, Helson L. The Kynurenine Pathway: A Primary Resistance Mechanism in Patients with Glioblastoma. Anticancer Res. 2017;37(5):2159–2171. doi: 10.21873/anticanres. 11551.

16. Matthew M, Neuberger A. Aminomalonate as an enzyme inhibitor. Biochem J. 1963;87(3):601–612.

17. Yang X, Palasuberniam P, Kraus D, Chen B. Aminolevulinic Acid-Based Tumor Detection and Therapy: Molecular Mechanisms and Strategies for Enhancement. Int J Mol Sci. 2015; 16(10):25865–25880. doi: 10.3390/ijms161025865.

18. Lawrence JE, Patel AS, Rovin RA, et al. Quantification of Protoporphyrin IX Accumulation in Glioblastoma Cells: A New Technique. ISRN Surg. 2014;2014:405360. doi:10.1155/2014/405360.

19. Nakamizo S, Sasayama T, Shinohara M, et al. GC/MS-based metabolomic analysis of cerebrospinal fluid (CSF) from glioma patients. J Neurooncol. 2013; 113(1):65–74. doi: 10.1007/s11060-013-1090-x.

20. Papait R, Magrassi L, Rigamonti D, Cattaneo E. Temozolomide and carmustine cause large-scale heterochromatin reorganization in glioma cells. Biochem Biophys Res Commun. 2009;379(2):434–439. doi:10.1016/j.bbrc.2008.12.091.

21. Happold C, Roth P, Wick W, et al. Distinct molecular mechanisms of acquired resistance to temozolomide in glioblastoma cells. J Neurochem. 2012;122(2):444–455. doi: 10.1111/j.1471-4159.2012.07781.x.

22. Kume K, Cantwell H, Neumann FR, Jones AW, Snijders AP, Nurse P. A systematic genomic screen implicates nucleocytoplasmic transport and membrane growth in nuclear size control. Hopper AK, ed. PLoS Genet. 2017; 13(5):e1006767. doi: 10.1371/journal.pgen. 1006767.

23. Liang Q-C, Xiong H, Zhao Z-W, et al. Inhibition of transcription factor STAT5b suppresses proliferation, induces G1 cell cycle arrest and reduces tumor cell invasion in human glioblastoma multiforme cells. Cancer Letters. 2009;273(1): 164–171. doi:10.1016/j.canlet.2008.08.011.

24. Giese A, Bjerkvig R, Berens ME, Westphal M. Cost of migration: invasion of malignant gliomas and implications for treatment. J Clin Oncol. 2003;21(8):1624–1636. doi: 10. 1200/JCO.2003.05.063.

25. Yeom S-Y, Nam D-H, Park C. RRAD promotes EGFR-mediated STAT3 activation and induces temozolomide resistance of malignant glioblastoma. Mol Cancer Ther. 2014; 13(12):3049–3061. doi:10.1158/1535-7163.MCT-14-0244.

26. Westphal M, Maire CL, Lamszus K. EGFR as a Target for Glioblastoma Treatment: An Unfulfilled Promise. CNS Drugs. 2017;31(9):723–735. doi: 10.1007/s40263-017-0456-6.

27. Garnier D, Meehan B, Kislinger T, et al. Divergent evolution of temozolomide resistance in glioblastoma stem cells is reflected in extracellular vesicles and coupled with radiosensitization. Neuro-oncology. July 2017. doi: 10.1093/neuonc/nox142.

28. Schimanski A, Ebbert L, Sabel MC, et al. Human glioblastoma stem-like cells accumulate protoporphyrin IX when subjected to exogenous 5-aminolaevulinic acid, rendering them sensitive to photodynamic treatment. J Photochem Photobiol B, Biol. 2016;163:203–210. doi:10.1016/j.jphotobiol.2016.08.043.

29. Stummer W, Novotny A, Stepp H, Goetz C, Bise K, Reulen HJ. Fluorescence-guided resection of glioblastoma multiforme by using 5-aminolevulinic acid-induced porphyrins: a prospective study in 52 consecutive patients. J Neurosurg. 2000;93(6): 1003–1013. doi: 10.3171/jns.2000.93.6.1003.

30. M A, H L M B. The AGT cytogenetics laboratory manual, 4th edition. Wiley-Blackwell. 2017.

31. Haddad BR, Gu L, Mirtti T, et al. STAT5A/B Gene Locus Undergoes Amplification during Human Prostate Cancer Progression. The American Journal of Pathology. 2013; 182(6):2264–2275. doi:10.1016/j.ajpath.2013.02.044.

32. Haddad B, Pabón-Peña CR, Young H, Sun WH. Assignment1 of STAT1 to human chromosome 2q32 by FISH and radiation hybrids. Cytogenet Cell Genet. 1998;83(1-2):58–59. doi: 10.1159/000015126.

33. KJ C, KA N, LM S, N A, JD R, BR H. Glutathione S-transferase pi amplification is associated with cisplatin resistance in head and neck squamous cell carcinoma cell lines and primary tumors. Cancer Res. 2003;1(63(23)):8097–8102.

34. Schindelin J, Arganda-Carreras I, Frise E, et al. Fiji: an open-source platform for biological-image analysis. Nat Methods. 2012;9(7):676–682. doi: 10.1038/nmeth.2019.

35. Doube M, Kłosowski MM, Arganda-Carreras I, et al. BoneJ: Free and extensible bone image analysis in ImageJ. Bone. 2010;47(6):1076–1079. doi: 10.1016/j.bone.2010.08.023.

36. Smith CA, Want EJ, O'Maille G, Abagyan R, Siuzdak G. XCMS: processing mass spectrometry data for metabolite profiling using nonlinear peak alignment, matching, and identification. Anal Chem. 2006;78(3):779–787. doi: 10.1021/ac051437y.

37. Ihaka R, Gentleman R. R: A Language for Data Analysis and Graphics. Journal of Computational and Graphical Statistics. 1996;5(3):299. doi:10.2307/1390807.

38. Pannkuk EL, Laiakis EC, Authier S, Wong K, Fornace AJ. Gas Chromatography/Mass Spectrometry Metabolomics of Urine and Serum from Nonhuman Primates Exposed to Ionizing Radiation: Impacts on the Tricarboxylic Acid Cycle and Protein Metabolism. J Proteome Res. 2017;16(5):2091–2100. doi: 10.1021/acs.jproteome.7b00064.

39. Mak TD, Laiakis EC, Goudarzi M, Fornace AJ. MetaboLyzer: a novel statistical workflow for analyzing Postprocessed LC-MS metabolomics data. Anal Chem. 2014;86(1):506–513. doi: 10.1021/ac402477z.

